# Analysis of microbial community structure of pit mud for Chinese strong-flavor liquor fermentation using next generation DNA sequencing of full-length 16S rRNA

**DOI:** 10.1101/380949

**Authors:** Zuolin Liu, Ying Han, Junwei Li, Runjie Cao, Hongkui He, Anjun Li, Zhizhou Zhang

**Author notes:** Equal contribution (Zuolin Liu, Ying Han). Contact: Zhizhou Zhang, PhD Prof 86-631-5683176.

## Abstract

The pit is the necessary bioreactor for brewing process of Chinese strong-flavor liquor. Pit mud in pits contains a large number of microorganisms and is a complex ecosystem. The analysis of bacterial flora in pit mud is of great significance to understand liquor fermentation mechanisms. To overcome taxonomic limitations of short reads in 16S rRNA variable region sequencing, we used high-throughput DNA sequencing of near full-length 16S rRNA gene to analyze microbial compositions of different types of pit mud that produce different qualities of strong-flavor liquor. The results showed that the main species in pit mud were *Pseudomonas extremaustralis* 14-3, *Pseudomonas veronii, Serratia marcescens* WW4, and *Clostridium leptum* in *Ruminiclostridium.* The microbial diversity of pit mud with different quality was significantly different. From poor to good quality of pit mud (thus the quality of liquor), the relative abundances of *Ruminiclostridium* and *Syntrophomonas* in Firmicutes was increased, and the relative abundance of *Olsenella* in Actinobacteria also increased, but the relative abundances of *Pseudomonas* and *Serratia* in Proteobacteria were decreased. The surprising findings of this study include that the diversity of intermediate level quality of N pit mud was the lowest, and the diversity levels of high quality pit mud G and poor quality pit mud B were similar. Correlation analysis showed that there were high positive correlations (r > 0.8) among different microbial groups in the flora. Based on the analysis of the microbial structures of pit mud in different quality, the good quality pit mud has a higher microbial diversity, but how this higher diversity and differential microbial compositions contribute to better quality of liquor fermentation remains obscure.

## 1. Introduction

Chinese strong-flavor liquor (CSFL) has a special status in the history of brewing because of its unique brewing process and flavor^1^. Strong-flavor liquor is produced using a cyclic solid fermentation process^2–3^. The fermentation pit is the infrastructure of strong-flavor liquor. It is built by digging a rectangular hole (about 2m × 3m × 2m) in the ground, and its inner wall is covered with a layer (of about 10cm thickness) of pit mud^4–5^. Mixture of the grain material of sorghum, wheat and corn together with Daqu (the fermentation starter) is put into the pit and sealed with common soil to conduct solid fermentation for 60-90 days^6–7^. After that, the fermented grains were taken out of the pit and distilled to produce strong-flavor liquor with ethyl caproate (1.5-3.0g/L) like pineapple and banana flavor^8^. Then the remaining fermented grains of the previous round are mixed with new raw materials, and inoculated with Daqu again for the next round of fermentation to repeat the above process periodically^9^. During the fermentation process, microorganisms in Daqu and pit mud play a key role. Daqu is a traditional starter made up of fungi and yeast, which is responsible for the simultaneous saccharification of raw materials and the fermentation of alcohol^7,10^. Pit mud provides a suitable habitat for a large number of microorganisms that participate in various biological processes, and their metabolites are often the precursors of many aroma components and various enzymes that promote the synthesis of aroma components^3,11,12^. Therefore, the composition of microbial communities in pit mud largely determines the flavor and quality of strong-flavor liquor.

In recent years, pit mud has attracted much attention due to its close relationship with the flavor and quality of Chinese strong-flavor liquor^5^. Yue et al.^13^ used anaerobic culture methods to screen and isolate bacteria in pit mud, mostly facultative anaerobic bacteria, and found *Clostridium* bacteria account for only about 1%. The results obtained are quite different from those obtained by uncultured technology, because only 0.1%-10% microbes in natural environment can be cultured and separated^14^. Zhao et al.^15^ used the PLFA method to detect the changing rules of microbial fatty acid in different years of pit mud in order to establish some correlation between fatty acid profile and pit mud quality. Zheng et al.^11^ reported that, by PCR-DGGE, the difference of gram positive (G+) bacteria, gram-negative (G^-^) bacteria and aerobic actinomycetes in the pit mud of different years was not significant. The banding information in the DGGE map may underestimate the diversity of dominant microorganisms in environmental samples and this method seems not suitable for sequences with PCR product fragment lengths greater than 500 bp. Hu et al.^5^ compared microbial compositions of anaerobic bacteria in a quality gradient of pit mud of a type of stong-flavor liquor by means of V4 16S rDNA high-throughput sequencing. Hong et al.^16^ combined V3-V4 16S rDNA high-throughput sequencing and metagenomic high-throughput sequencing to analyze the microbial compositional dynamics during wine-making and discovered the important role of *Lactobacillus brevis* in rice wine fermentation. However, the current second-generation high-throughput sequencing is limited in read length and cannot measure full-length of ribosome RNA gene; the longest length can only measure 1-2 variable regions^17^ and analysis of microbial communities thus revealed significantly lower species abundance than that obtained using full-length sequences^18^.

The role of pit mud in the fermentation of liquor is mainly reflected in its microbial flora. After the long-term brewing environment of domestication and selection of pit mud microorganisms, and the flora of the pit mud gradually becomes complicated and form a specific dominant groups of bacteria^19–20^. To overcome the limitations brought by short sequencing reads in the microbial composition analysis at the species level due to the high similarity of short 16S rRNA variable regions^21^, we used (near) full-length 16S rRNA sequencing to analyze microbial community of pit mud with different quality in GujingTribute liquor (also strong-flavor liquor). Using different statistical analysis methods to analyze different quality bacterial community of pit mud by R (https://www.r-project.org/), the authors tried to find the key strains that may most affect liquor quality, and accumulate more theoretical knowledge on how to improve the quality of strong-flavor liquor.

## 2. Materials and methods

### 2.1. Sample collection

Pit mud samples were taken based on liquor qualities from the pits in which the liquor is fermented and distilled. In general, the older the pit, the better the liquor. However, not all old pits produce high quality liquor during some period, and not all young pits produce normal or bad quality of liquor during some period. So we randomly sampled from 19 pits of GujingTribute liquor (Bozhou, China) only based on the liquor quality (**<u>G</u>**ood, **<u>N</u>**ormal and **<u>B</u>**ad). The liquor quality was judged and scored (Table 1s) according to GujingTribute production standard of strong-flavor liquor. Samples of pit mud were taken using the nine-point sampling method (Figure 1s). Nine red spots clearly indicated the specific locations of the sampling of pit mud: the central point of four sidewalls, the four corners of the pit bottom and the center of the pit bottom. A 2cm×2cm×2cm mud mass at each of the nine points was picked, and then equally mix the nine points of mud mass to get one pit mud sample.

### 2.2. DNA extraction of samples

The sample DNA was extracted by using Sangon Biotech (Shanghai, China) columnar soil genomic extraction kit (SK8263). Due to the protective effect of starch sticky substances in pit mud or the majority of Gram-positive bacteria, the genome was difficult to extract. Therefore, this experiment slightly modified the first step of the kit experimental procedure: Weigh 100-300 mg of soil, add 400 μl of pre-heated Buffer SCL at 65°C, and then place it in -80°C for 15 min and a water bath at 90°C. 5min repeated freezing and thawing three times to promote the full lysis of microorganisms, and then vortex mixing, placed in a 65 °C water bath for 5min, and then in strict accordance with the steps of the kit extraction. The obtained DNA solution was stored at -20°C for further use.

### 2.3. 16S rDNA amplification

The 16S rDNA was used as a template for PCR amplification using universal primers 27F (5’-AGA GTT TGA TCM TGG CTC AG) and 1492R (5’-TAC GGY TAC CTT GTT ACG ACTT) by NPK02 kit (Weihai XiaoDong Biotech). The 12 μL PCR system was used, 6 μL 2×NPK02 buffer, 0.8 μL (2 μM) upstream primers, 0.8 μL (2 μM) downstream primers, 0.2 μL Taq enzyme,1 μL (10ng/μL) genome template and 3.2 μL ddH_2_O to 12 μL. The PCR amplification procedure was: Preheat at 94°C for 5 min, amplify for 30 cycles (94°C-30s, 60°C-30s, 72°C-40s), plus the extension at 72°C for 5 min. The PCR production was subjected to agarose gel electrophoresis (Figure 2s) and the target PCR product was purified.

### 2.4 Full-length 16S rRNA sequencing

Single molecule sequencing of full-length rDNA was performed using the Pacbio Sequel platform assigned to Personalbio (Shanghai, China).

### 2.5 Sequence data analysis

First, the questioning sequence was identified using QIIME software (Quantitative Insights Into Microbial Ecology, v1.8.0, http://qiime.org/)^22^. In addition to requiring a sequence length of ≥500 bp and not allowing the presence of a fuzzy base N, we also removed: 1) sequences with 5’ primer mismatch bases >5, and 2) sequences contain consecutive same bases >8. Subsequently, USEARCH (v5.2.236, http://www.drive5.com/usearch/) was called by the QIIME software (v1.8.0, http://qiime.org/) to check and reject the chimeric sequence. Using QIIME software, UCLUST, a sequence comparison tool^23^, was invoked to perform the merging and OTU partitioning on the sequence obtained above with a sequence similarity of 97%, and the most abundant sequence in each OTU was selected as the representative sequence of the OTU. For each OTU representative sequence, the default parameters are used in the QIIME software. The OTU representative sequence is compared with the template sequence of the Greengenes database to obtain the taxonomic information corresponding to each OTU. In order to explain the bacterial richness and evenness, QIIME was used to calculate the diversity index including Chao1, Shannon, Simpson and ACE parameters. Statistics and correlation analysis were performed with R.

### 2.6 Nucleotide sequence accession number

The sequencing data are available at the DDBJ database under accession no. PRJDB7224.

## 3. Results and Discussion

### 3.1. Analysis of full-length 16S rRNA gene sequencing of bacterial community in pit mud

In this study, 19 pit mud samples generated a total of 192,834 reads. After the reads was filtered and quality controlled, the average processed reads for each sample were 10149 (Table 1). In order to cluster the reads according to sequence similarity, we used the 97% similarity level to perform OUT (operational taxonomic unit) classification. The OTUs in each pit mud sample vary, with the largest number of OTUs found in the G3 samples (323 OTUs) in G pit mud, and the lowest number of OTUs found in the N6 samples (98 OTUs) in N pit mud. The OTUs of G pit mud are significantly higher than the N pit mud (Figure 1A). During the conversion from B to G pit mud, the OTUs could first decreases, then increases, due to the specific acclimatization process of the bacteria in the pits.

**Figure 1.**
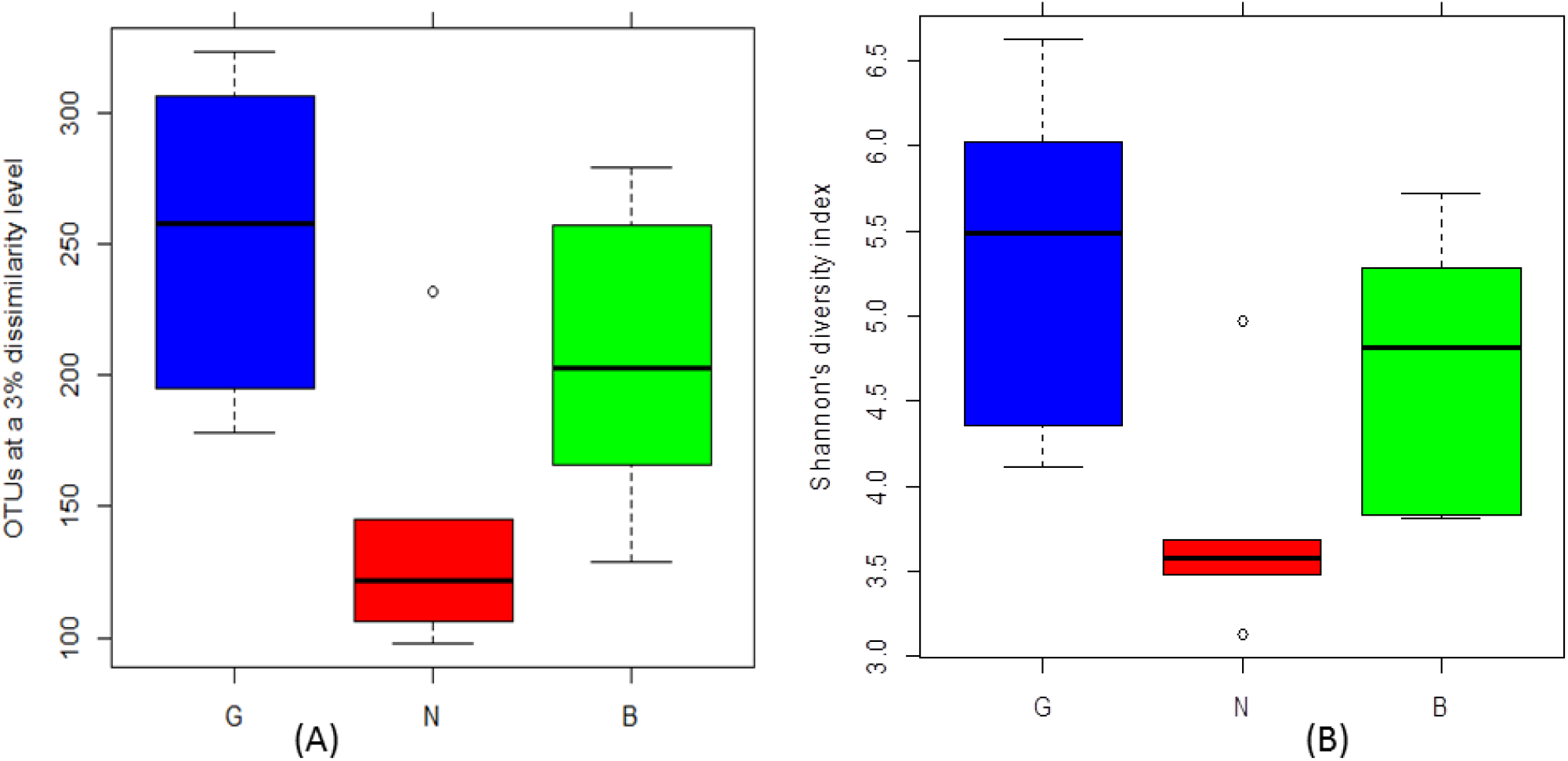
(A) OTU classification of bacterial in pit muds with different quality; (B) Shannon’s diversity index of different quality pit mud. In both (A) and (B), G and N had significant difference while B had no significant difference from either G or N (by Tukey HSD p<0.05).

### 3.2. Analysis of alpha diversity of bacterial community in pit mud

The use of alpha diversity in the analysis of the number of microbial species in a single pit mud sample mainly refers to the richness and diversity of the microbial community. The statistics commonly used to estimate the richness of microbial communities are Chao1 and ACE. The commonly used estimates of microbial community diversity are Shannon and Simpson. The higher the diversity index, the higher the diversity of the community. The Shannon diversity index comprehensively considers the richness and evenness of the community, and can be used as a microbial indicator that responds quickly to the external environment and is widely used to evaluate soil, water quality, and fermentation performance of the fermentation system^24–26^. Pit muds of different quality have different degrees of diversity in the bacterial community, but in general the bacterial community in the G pit mud was more diverse than those in the B and N pit mud. In particular, there was a significant difference between G and N, or B and N, but there was no significant difference between B and G (or N) pit mud (Table 1 and Figure 1). This is consistent with the results of OTUs quantitative analysis.

**Table 1.**
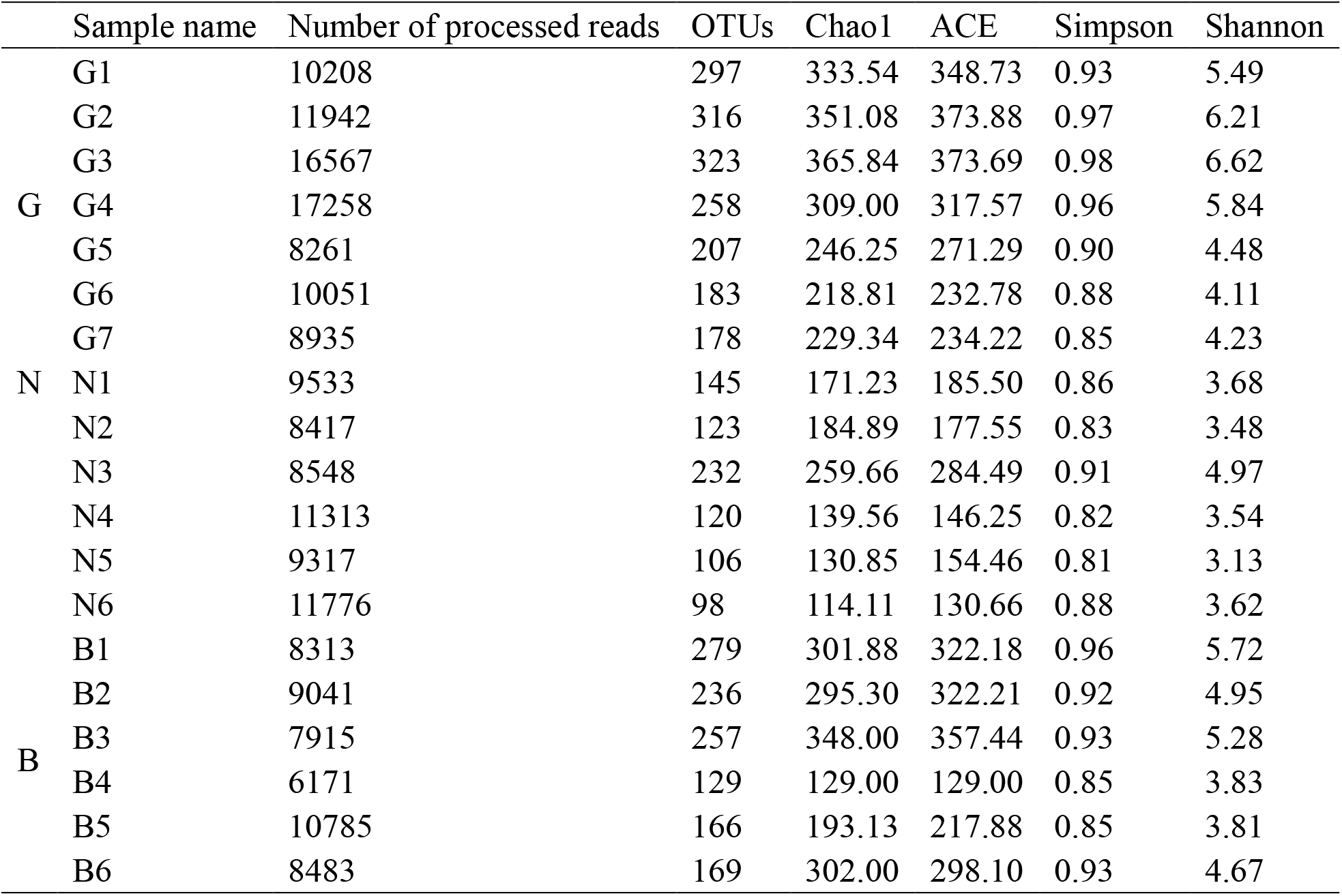
Number of processed sequences, clustered OTUs and alpha diversity estimates of bacterial communities in each sample

Shannon-Wiener curves^27^ can be used to characterize the microbial community diversity and richness of pit mud samples at different sequencing depths. When the curve is flattened, it means that the sequencing amount can cover all species in the sample and can represent the vast majority of microbial information in the sample. According to Shannon-Wiener curves of bacterial community in pit muds with different years (Figure 3 s), with the increase of the amount of sequencing, the trend of increasing the diversity index gradually slowed down and eventually reached saturation. This can fully demonstrate that the current reads amount is reasonable, and the diversity of bacterial communities in pit mud can be basically detected.

### 3.3 Analysis of bacterial communities in pit muds at the phylum level

The OTU representative sequences were compared with the Greengene database. Fifteen phyla were used to classify microbial communities in the pit mud (Figure 2A). The diversity of G pit mud at phylum level was significantly higher than that of B and N. The above OTU analysis and Shannon’s diversity index analysis were consistent with each other. Surprisingly, the microbial diversity of N pit mud was the lowest in general among different qualities of pit mud. The potential reason for this may be that in the complex pit environment, the N pit mud is in a transitional state where the flora evolves, and some part of the flora not adapted to environmental factors is eliminated. But like Proteobacteria, which are likely adapted to the environment, are becoming more and more abundant. With the increase of the liquor quality, the microbial community’s flora diversity has gradually increased and is higher than that of the lower quality pit mud. Firmicutes and Proteobacteria occupy the absolute dominant position in different types of pit mud, and their sum of relative abundance was about 89.6%. Therefore, the bacteria in these two phyla play a greater role in the stability of pit mud. Pit mud bacteria were mainly distributed in Firmicutes, Proteobacteria, Bacteroidetes and Synergistetes. Ding et al.^28^ found that the main phylum distribution of pit mud in three different years (between 2 to 30 years) was also the above four phyla. We conducted a significant difference analysis of the relative abundances of 15 phyla in the pit mud (Figure 2B). The G and N pit mud were significantly different in Proteobacteria, Choroflexi, Cloacimonetes, and Euryarchaeota, while the N and B pit mud were significant different in Actinobacteria. However, the G and B pit mud did not have significant differences on any phylum except for Proteobacteria and Firmicutes. As the liquor quality increases, the relative abundance of Firmicutes increased, and the relative abundance of Proteobacteria between N and B was also increasing. However, Proteobacteria dropped to about 29.14% during N conversion to G. The relative abundance of Bacteroidetes^29^ and Synergistetes also increased in the B conversion to G; while the relative abundance of Actinobacteria was reduced. The relative abundance of Euryarchaeota is 0.04%, and Euryarchaeota in the G pit mud is 4.45 times that of B pit mud. Therefore, microorganisms between the four phyla of Firmicutes, Euryarchaeota, Bacteroidetes and Proteobacteria may have a somewhat similar niche and there may be a symbiotic relationship between these microorganisms^30–32^. For example, Vanwonterghem et al^33^. found that Clostridiale, Syntrophomonas, Bacteroidales, Syntrophobacterales, Methanomicrobiales, and Methanosarcinales in the above four phyla cooperated to complete the anaerobic digestion process, starting the complex carbon source cellulose to produce organic acids and hydrogen, and ultimately producing methane.

**Figure 2.**
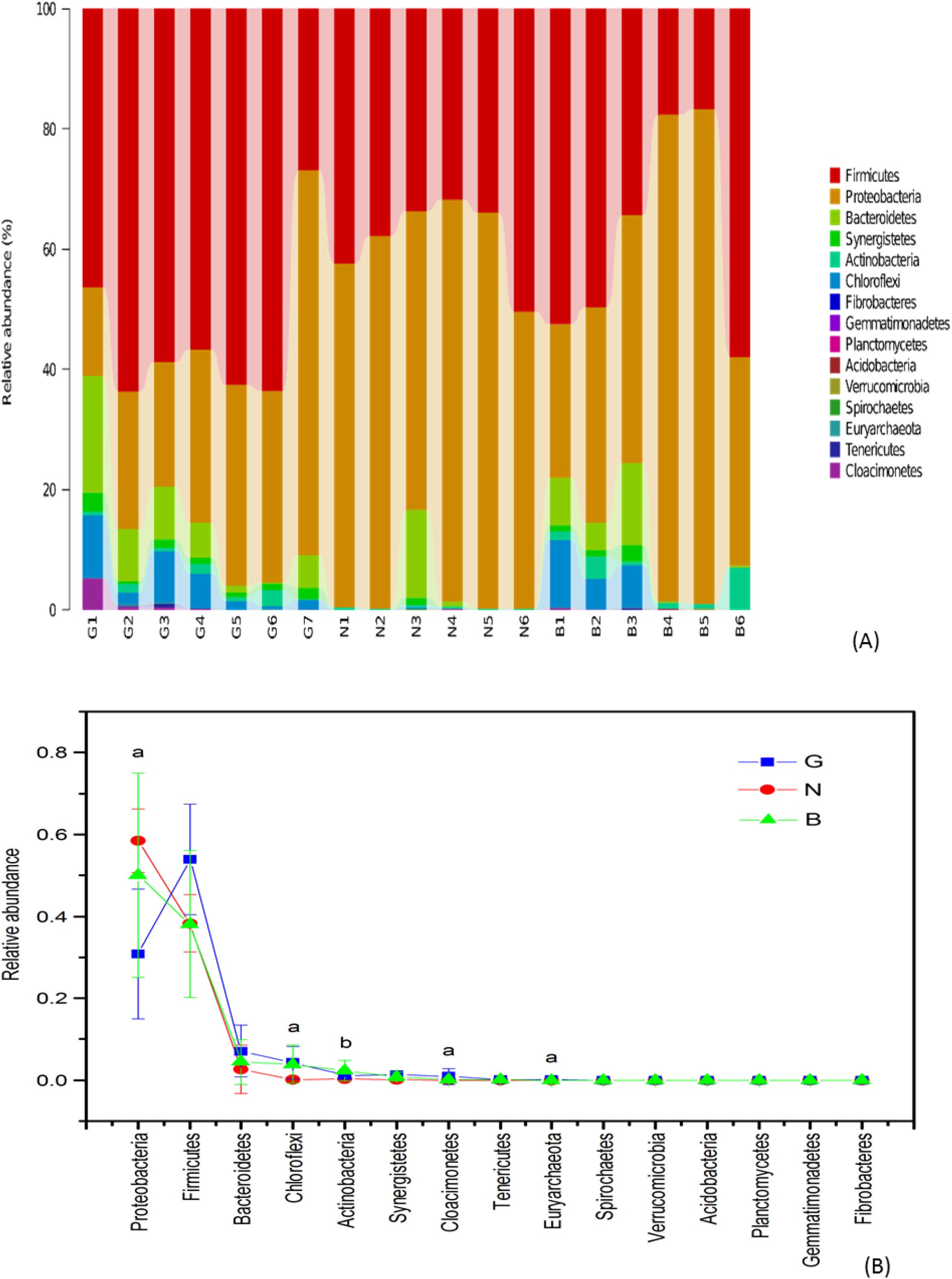
(A) Distribution of bacterial community in three types of pit muds at phylum level. (B) Relative abundance difference of 15 phyla in three types of pit muds. *(a* indicating a significant difference of that phylum between G and N and *b* indicating a significant difference of that phylum between N and B by ANOVA, p < 0.05).

### 3.4 Analysis of bacterial communities in pit muds at the genus level

After comparison with the Greengene database, the community in the pit mud was classified into 190 genera. The two genera, *Pseudomonas* and *Ruminiclostridium,* occupied an absolute dominant position in pit mud, and the sum of their relative abundance was about 54.3% (Figure 3A). There were 30 genera in the entire samples whose relative abundances were all greater than 0.1%, and the total abundance of these 30 genera accounts for 93.9% of the microbial community. These 30 genera belonged to Proteobacteria, Firmicutes, Bacteroidetes, Chloroflexi, Actinobacteria, Synerqistetes, and Cloacimonetes. These 7 phyla were just the most abundant ones in the pit mud, in which Firmicutes contained 15 genera. The clustering results of 30 genera (Figure 4s) found that the 19 samples were clearly divided into two categories. The 8 samples (the left part of Figure 4s) were clustered into one category, which contained more samples of pit mud B, and the 11 samples (The right part of Figure 4s) were clustered into another category, which mainly contained G pit mud samples, while the B pit mud samples are scattered in the two categories. This phenomenon may help to explain that the inner differences in the pit mud of G and N within each group were smaller, while the differences within samples were larger in the B group. We then analyzed 13 of the 30 genus (genus with a total relative abundance greater than 1%). The relative abundance of these 13 genera accounted for 88.0% of the total abundance. With the growth of liquor quality (Figure 3B), the relative abundance of *Pseudomonas* appeared to have increased first and then decreased, and its abundance in the pit mud G was lower than pit mud of B. However, contrary to *Pseudomonas*, *Ruminiclostridium* appeared to have less relative abundance and then increased, and its abundance in pit mud G was higher than pit mud B. In particular, the relative abundance of *Leptolinea* and *Ercella* decreased in pit mud during quality conversion of B to N, and these two genera increased in the pit mud from the quality N conversion to G. Interestingly, the total abundance of *Lactobacillus* was very close to the sum abundance of *Lactobacillus*, *Leptolinea, Levilinea, Olsenella and Petrimonas* either in B or G, suggesting that *Lactobacillus* increased from B to N, and then decreased from N to G. Indeed, *Lactobacillus*, *Ruminococcus, Calormator, Clostridium, Syntrophomonas*, etc. play important roles in the pit mud of strong-flavor liquor^34^. *Lactobacillus* can produce carbohydrates and other metabolites by fermenting carbohydrates, and its rapid increase will harm the stability of the bacterial structure in pit mud. Its relative abundance has gradually increased from pit mud B to N. *Clostridium* as a fermentation acid producing microorganisms, its fermentation products are mainly butyric acid, caproic acid, alcohol, CO_2_ and H_2_ ^35–36^. Its relative abundance has gradually decreased during the time in which the quality of liquor is improved to high quality. *Syntrophomonas* can degrade long-chain fatty acids into acetic acid and H2 ^37^. Its relative abundance has gradually increased when the quality of liquor is improved to high quality. The above research results show that the complexity of the microbial community in the pit mud of G was larger than the pit mud of B and N, which theoretically proves that the quality of pit mud gradually increases with the growth of the liquor quality.

**Figure 3.**
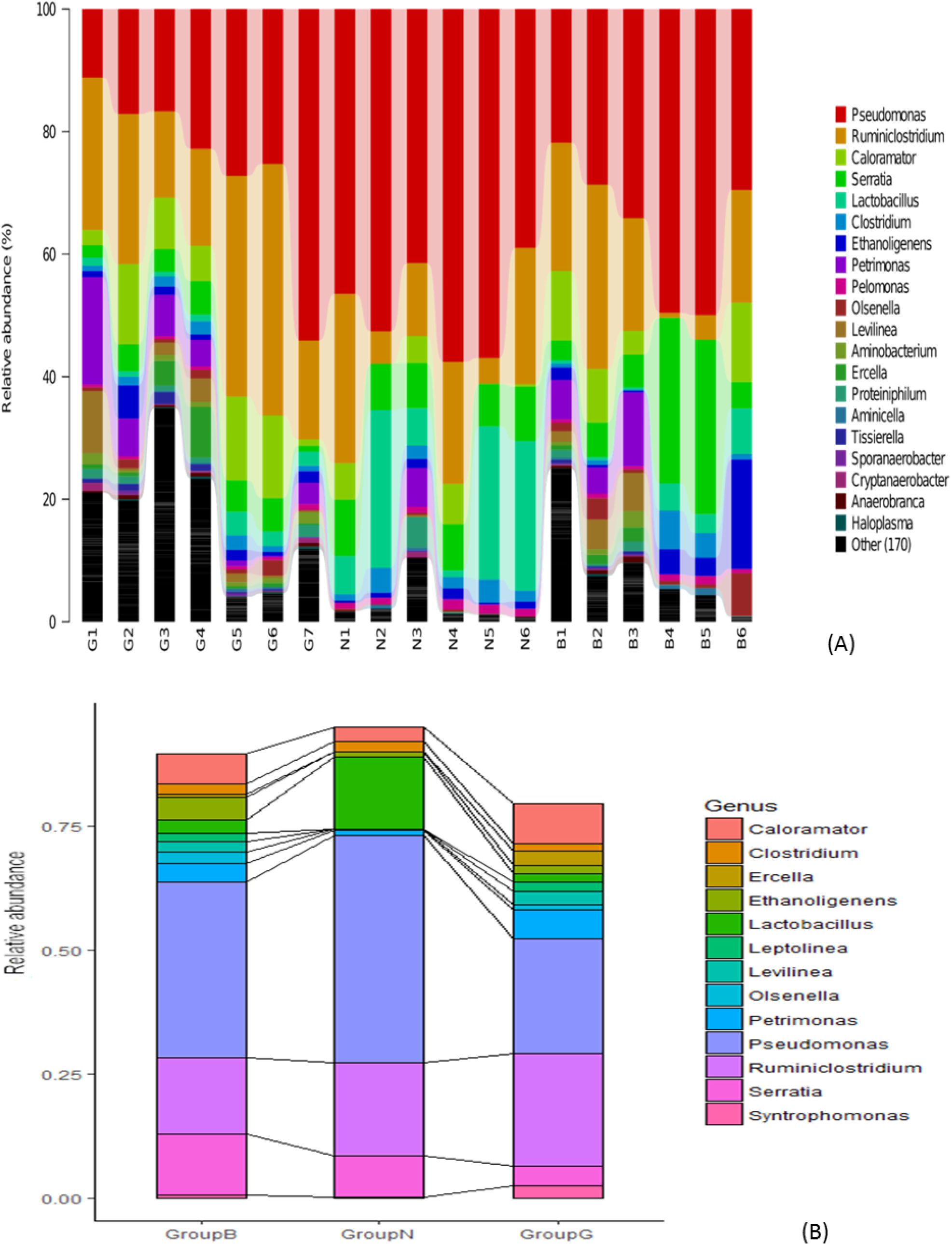
(A) Distribution of bacterial community in pit muds at genus level (the most dominant 20 genus); (B) Relative abundance changes of 13 genera (with relative abundance greater than 1%) in the pit mud over quality transformation.

### 3.5. Analysis of bacterial communities in pit muds at the species level

After comparison with the Greengene database, the bacterial community in the pit mud was attributed to 352 species (the most dominant 20 species shown in Figure 4). The diversity of the pit mud of G was significantly higher than that of the B and N pit mud. The N pit mud may be in the succession of the community, with a minimal diversity. *Pseudomonas extremaustralis* 14-3 (19.3%) and *Pseudomonas veronii* (15.7%) accounted for absolute superiority in the species level, and *Pseudomonas extremaustralis* 14-3 was closely related to *Pseudomonas veronii* with a similarity of 99.7%^38^. *Pseudomonas extremaustralis* 14-3 mainly produces polyhydroxybutyrate (PHB), a short chain length polyhydroxyalkanoate (sclPHA), which is not common in *Pseudomonas* species^39^. Both of these are substances that are easily degradable and can be converted into flavor molecules of strong-flavor liquor by other bacterial groups. Pit mud also contains *Clostridium leptum* (17.5%), which has not been accurately classified. It is also known as Clostridial cluster IV group, including *Faecalibacterium prausnitzii,* certain species of Eubacterium, and Ruminococcus^40^. *F. prausnitzii* was the most abundant member of the group and was known as the main producer of butyric acid through the fermentation of carbohydrates^41^. We conducted Pearson correlation analysis of the top 30 most abundant speciess in pit mud samples, including identified species and inaccurately classified groups (Figure 5A). The correlation coefficient |r|≥0.8 was highly correlated, and there were 19 highly positive correlations in pit mud samples (Figure 5B). It showed that in the same phylum, the highly relevant probability of occurrence is higher than between different phyla. *Pseudomanas extremaustralis* 14-3, *Pseudomonas veronii* and *Pelomonas saccharophila* had a high degree of positive correlations whit each other, and belong to Proteobacteria; every pit mud sample had these three bacteria, and the former two were all high abundance species in pit mud. These three also formed a highly positively related network with *Desulfotomaculum kuznetsovii* DSM 6115, *Oxobacter pfennigii, Leptolinea tardivitalis, Acetobacterium bakii* and *Syntrophomonas bryantii. Leptolinea tardivitalis* belongs to Chloroflexi, and the other six species belong to Firmicutes. *Syntrophomonas* is a kind of bacterium that lives in anaerobic environment and strictly depends on hydrogen-consuming metganogens to degrade butyric acid, and plays an important role in the anaerobic degradation of organic matter^42^. Although archaea such as methanogens are relatively low in pit mud, they can exert control over the pH of pit mud and have some effects on the degradation of harmful substances in pit mud. The interaction between methanogens and other microorganisms can not only be able to promote the formation of caproic acid, but also improves the stability of microbial communities in pit mud by reducing the pressure of hydrogen^43^. The methanogens have strict requirements for the growth environment. The 8 bacteria (Middle position of the Figure 5B) were estimated to have a mutually beneficial relationship with methanogens. The highly negative correlation among microbial species seemed not appear in the pit mud. Moreover, *Lactobacillus* in the pit mud is a concern, and the content of *Lactobacillus* is reduced during the process from common pit mud to high-quality pit mud^5^. *Lactobacillus acetotolerans* and *Lactobacillus acidipiscis* were negatively related to *Syntrophomonas erecta subsp. sporosyntropha* and many other species. In general, the *Lactobacillus* content of pit mud in pit mud of G was less than that in N and B pit mud, though the content of *Lactobacillus* itself is not enough to determine the quality of the pit mud. The main reason for the change in the quality of this pit mud may be that, in the process of liquor brewing, with the growth of pit age, the environmental factors in the pit mud will gradually change, thus the diversity of the prokaryotic microbial community in the pit mud will finally increase^34^.

**Figure 4.**
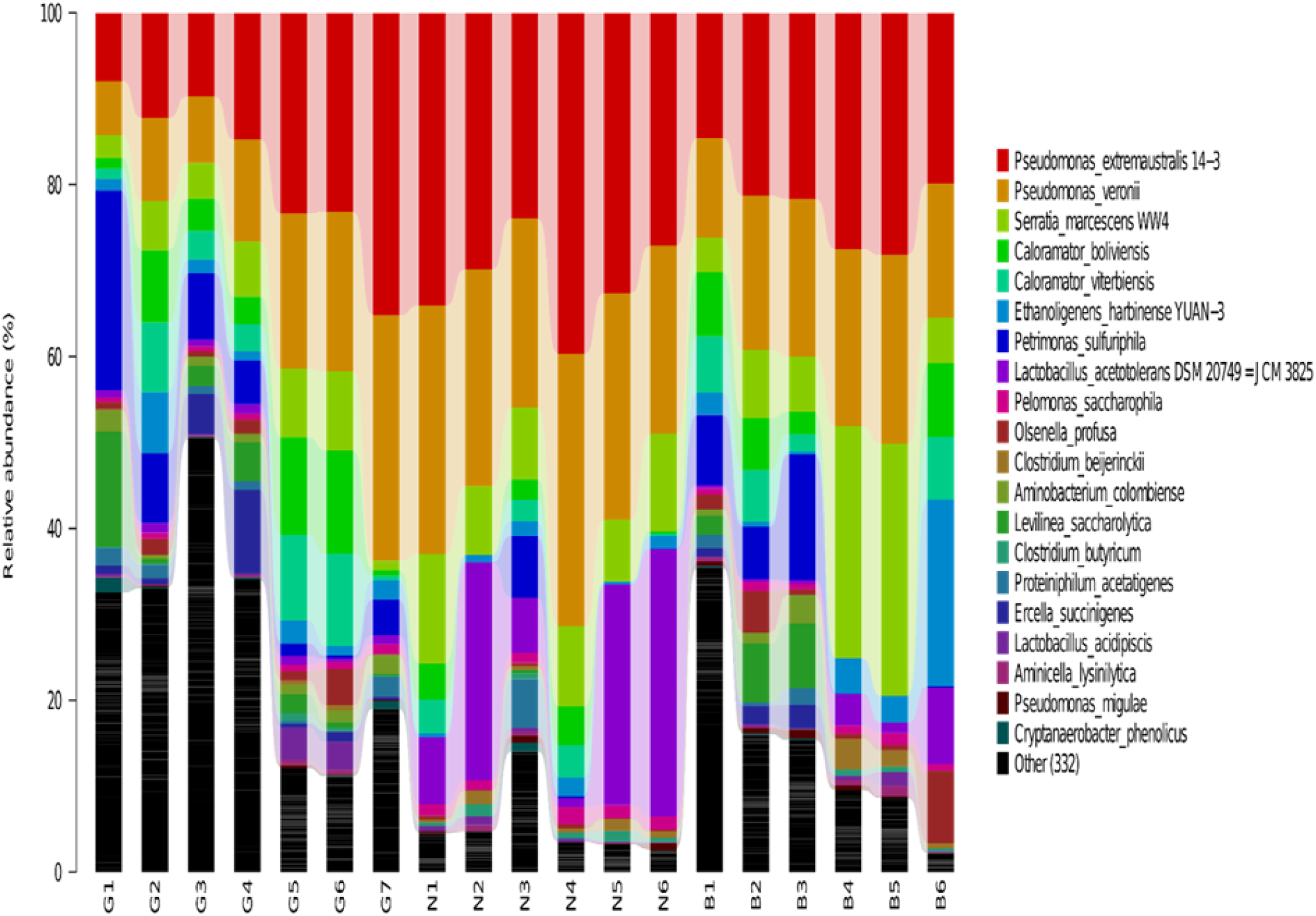
Distribution of bacterial community in pit muds at species level (the most dominant 20 species).

**Figure 5.**
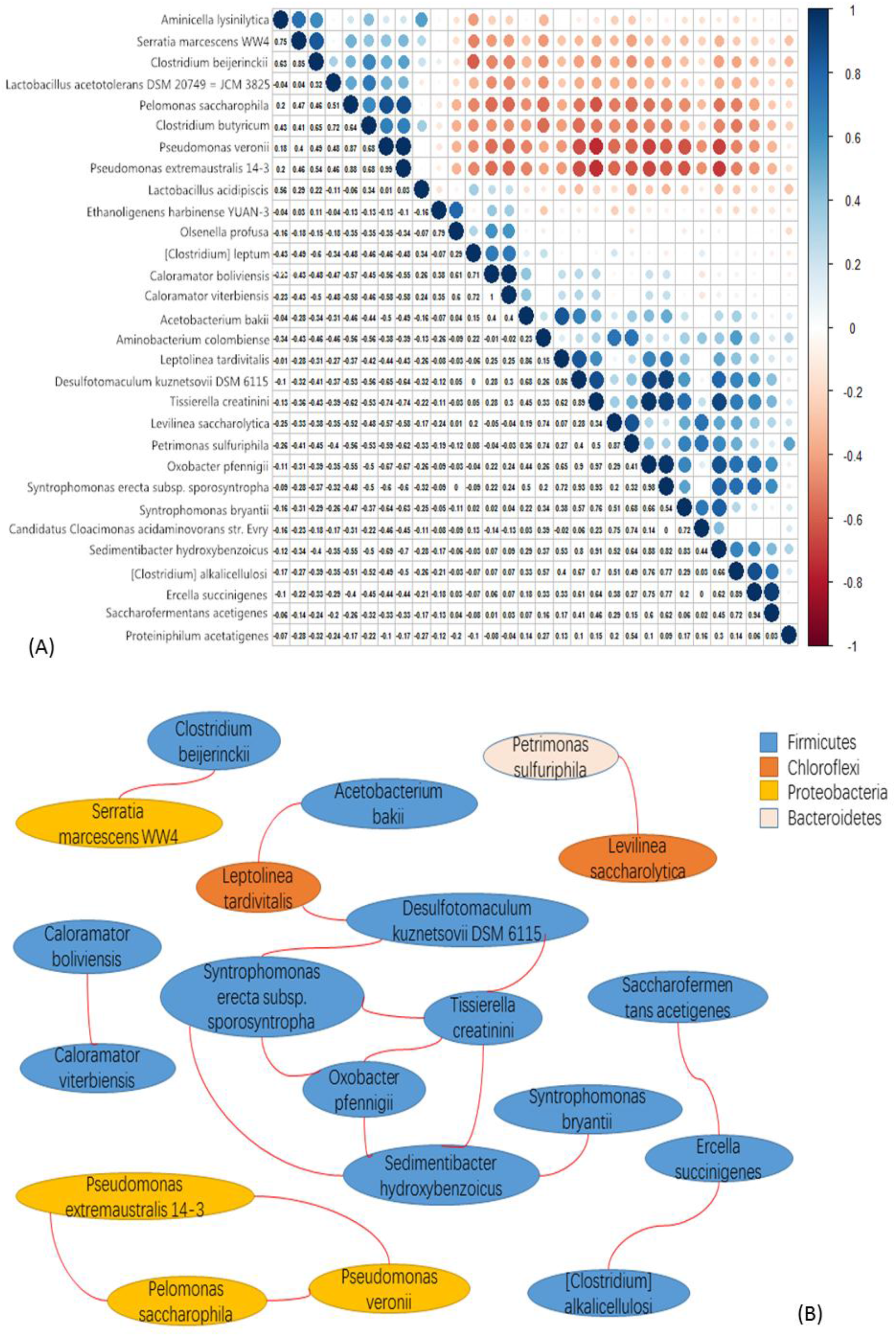
(A) Pearson analysis of the 30 most dominant classifications in pit mud samples, including identified species and inaccurately classified groups (r>0 indicates positive correlation and r<0 indicates negative correlation, |r|≥0.8 indicate highly correlated,0.5≤|r|<0.8 indicates moderate correlation, 0.3≤|r|<0.5 indicates low correlation and |r|<0.3 indicates very weak correlation); (B) Pearson correlation analysis depicts inter-species correlation in a highly relevant network diagram (r ≥0.8).

## 4. Conclusion

This study provided an approach to decipher the microbial composition and analyze its biodiversity by using full length 16S rRNA gene high throughput sequencing. Comparing with 16S variable region based next generation sequencing, this study would provide more species-level information that may be useful for functional considerations. However, pit mud microbial community is a very complex entity in which many species have been never found and studied, and some of their species-level taxonomic names may be not so accurate due to the lack of information in present databases.

In this study, the differences in microbial community structures between pit mud of different levels of fermentation quality were observed and analyzed. Biodiversity is the basis of the stability of the ecosystem. Higher levels of biodiversity in the pit mud are conducive to the function of the pit mud’s ecosystems. The biological diversity of the pit mud of B type in this study may be relatively close to that of ordinary soil. But G type of pit mud has an apparently higher diversity than N and B, with a surprising observation that N type of pit mud has the lowest level of microbial diversity. It is likely that high percentage of *Lactobacillus* in the N type of pit mud may contribute to the above surprising situation. In general, the quality of Chinese strong-flavor liquor has a certain relationship with the quality of the pit mud, and the microbial compositions between pit mud of different levels of fermentation did have differential characteristics as revealed in this study, but molecular mechanisms behind the above observations are waiting for more investigations.

## 5. Acknowledgements

This study was supported by the following funds: NSFC (No.31071170); GujingTribute fund (2016-1); GREDBIO (201401); the Key research and development plan of Shandong Province (grant number 2016GSF115022); and the Natural Science Foundation of Shandong Province (grant number ZR2018MC002).

## Supporting documents

**Table 1s.**
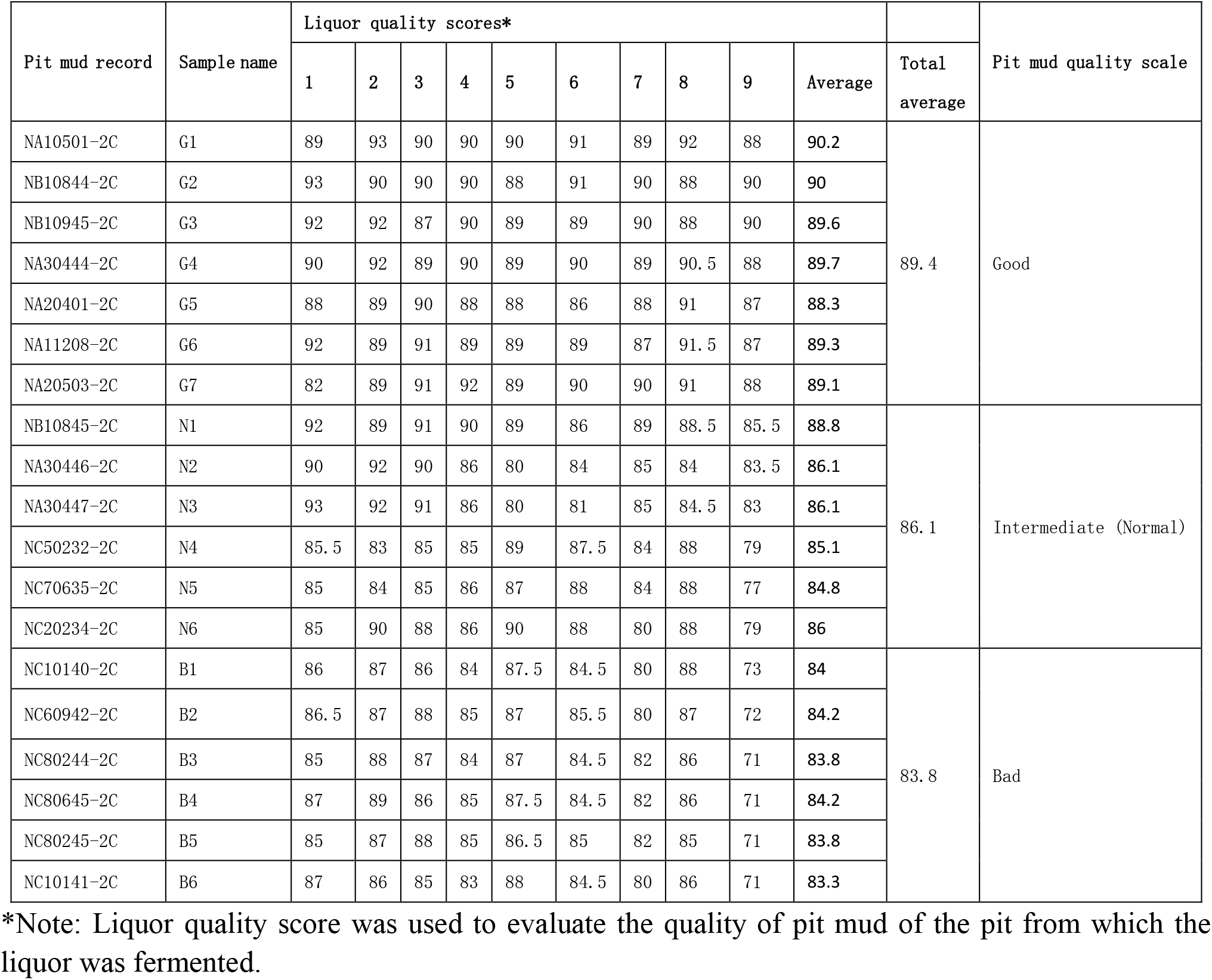
Sampling record and liquor quality scores

**Figure 1s.**
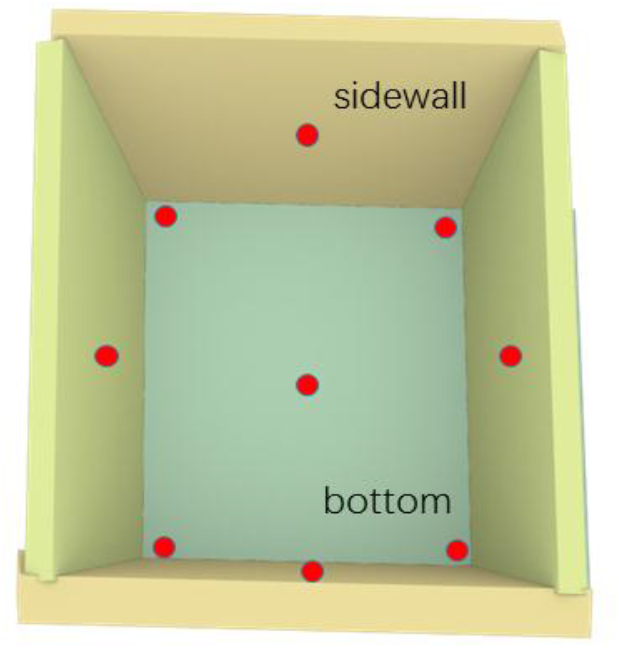
The nine spots sampling method for pit mud samples (the detailed sampling sites were shown above with nine red spots)

**Figure 2s.**
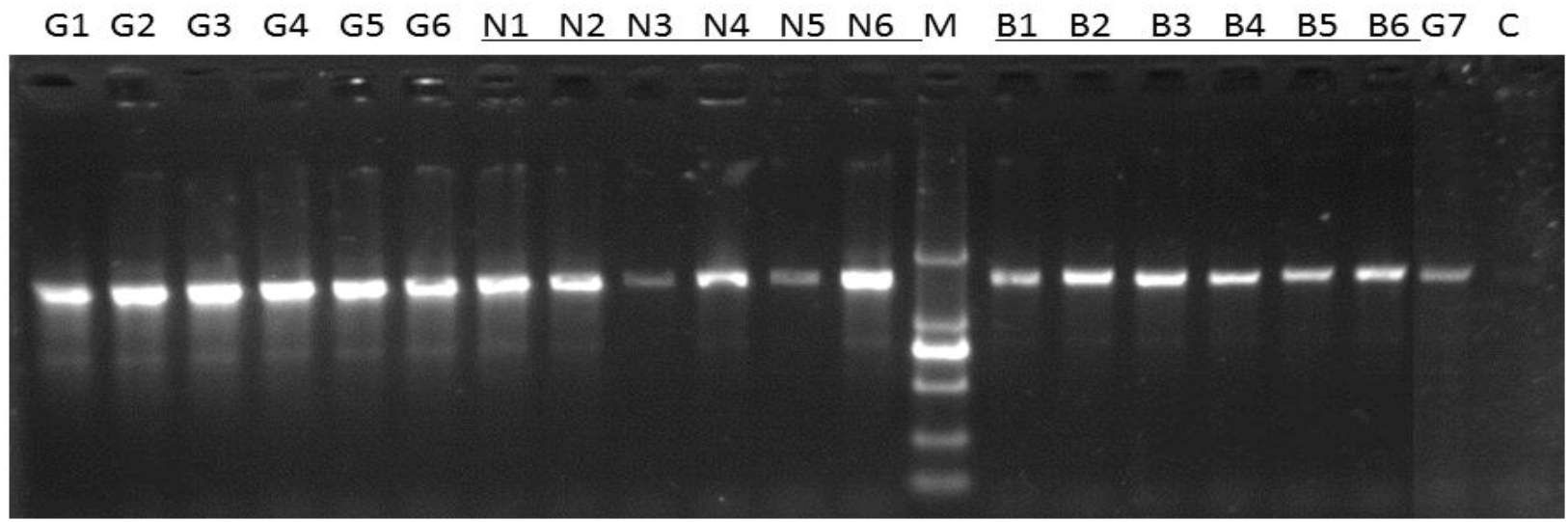
Amplification of 16S full length rDNA in 19 samples using NPK64 kit (Weihai Xiaodong Biotech). M: DNA marker DL2000 (2000, 1000, 750, 500, 250, 100bp); C: the blank control (amplification without the template DNA). PCR condition: 95°C -3min; (95°C -30s + 56°C -70s + 72°C -60s) ×37 cycles; 72°C -3min. 12ul PCR product was resolved in 1.2% agarose gel.

**Figure 3s.**
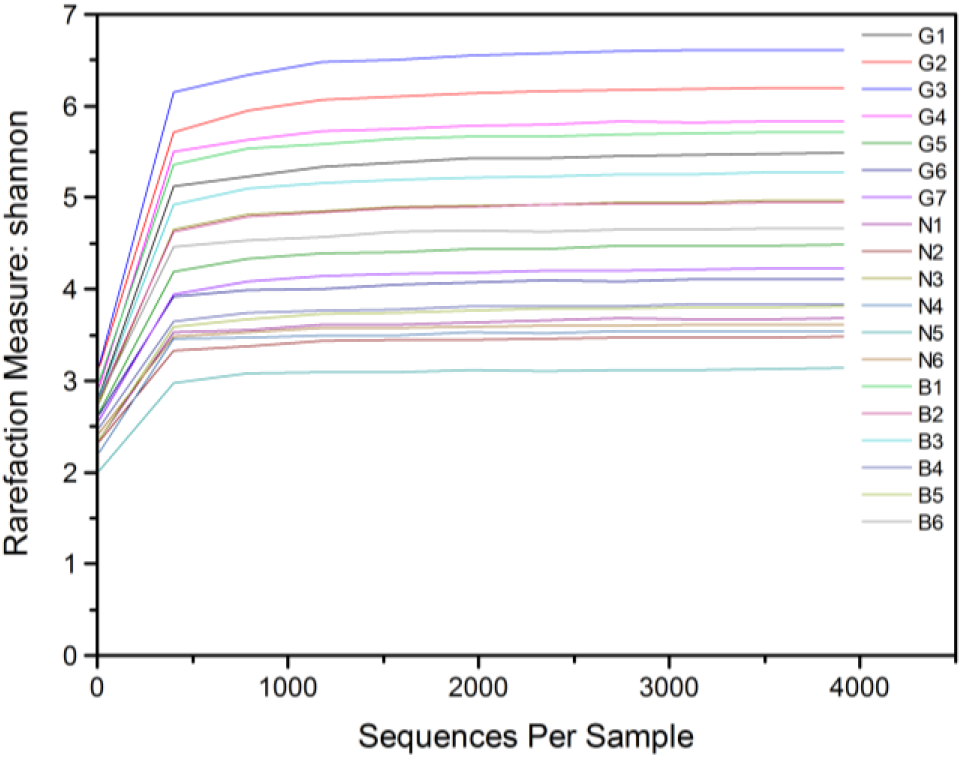
Shannon-Wiener curves of bacterial communities in pit muds.

**Figure 4s.**
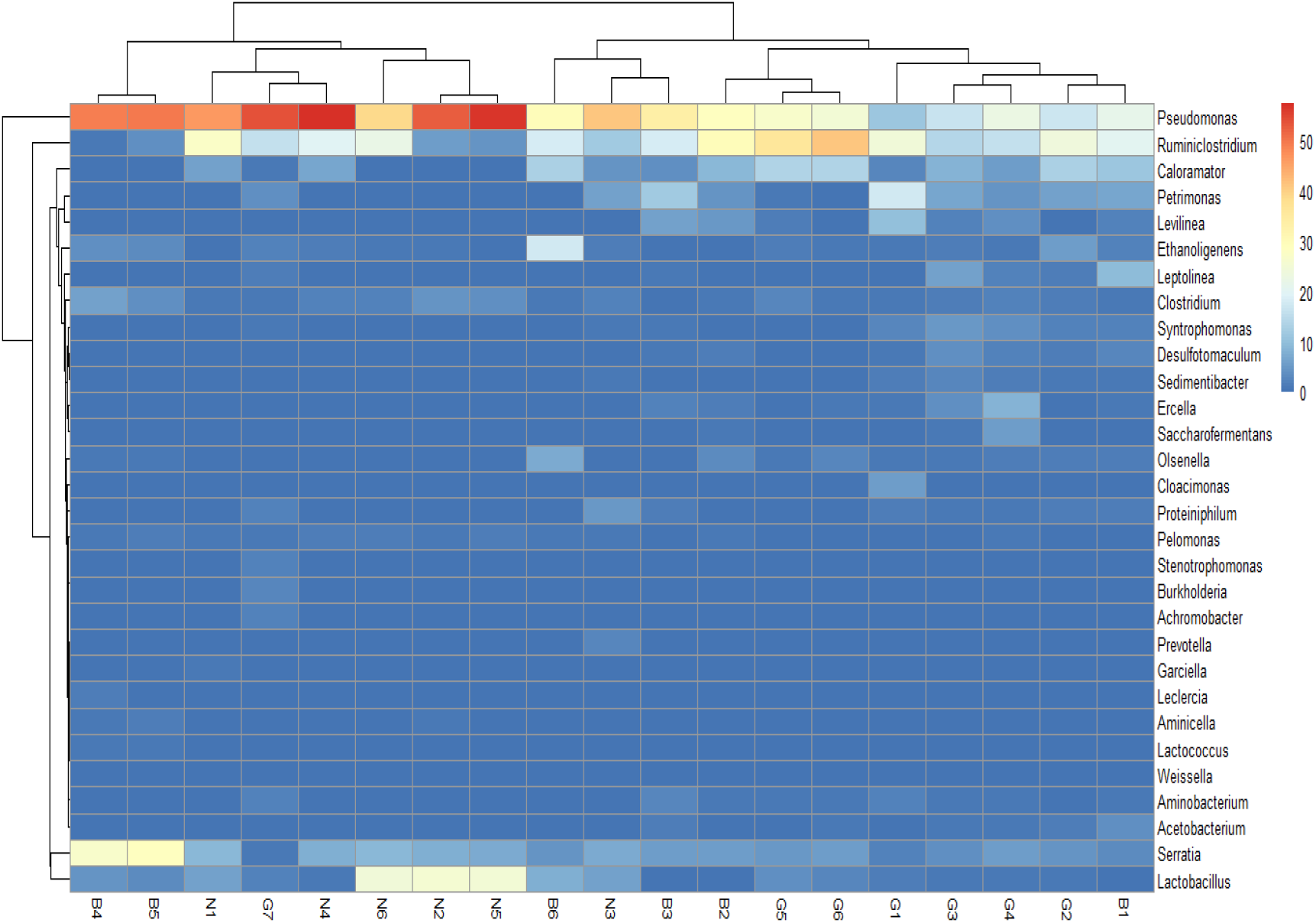
A dendrogram illustrating similarities among the bacterial profiles of different pit mud samples. The most dominant 10 genera of every sample led to a namelist of total 30 genera whose abundances were all greater than 0.1%, and the color scale indicates the relative abundance of each genus.

## References

1. Yue, Y. Y., Zhang, W. X., Rui, Y., Zhang, Q. S. & Liu, Z. H. (2007). Design and operation of an artificial pit for the fermentation of Chinese liquor. Journal of the Institute of Brewing, 113(4), 374–380.

2. Zhang, W. X., Qiao, Z. W., Shigematsu, T., Tang, Y. Q., Hu, C., Morimura, S. & Kida, K. (2015). Analysis of the bacterial community in zaopei during production of Chinese Luzhou – flavor liquor. Journal of the Institute of Brewing, 111 (2), 215–222.

3. Tao, Y., Li, J., Rui, J., Xu, Z., Zhou, Y., Hu, X., Wang, X., Liu, M., Li, D. & Li, X. (2014). Prokaryotic communities in pit mud from different-aged cellars used for the production of Chinese strong-flavored liquor. Appl Environ Microbiol, 80(7), 2254–2260.

4. Hu. X., Hai, D. & Yan, X. (2015). Identification and quantification of the caproic acid-producing bacterium *Clostridium kluyveri* in the fermentation of pit mud used for Chinese strong-aroma type liquor production. International Journal of Food Microbiology, 214, 116–122.

5. Hu, X., Du, H., Ren, C. & Xu, Y. (2016). Illuminating anaerobic microbial community and co-occurrence patterns across a quality gradient in Chinese liquor fermentation pit muds. Applied & Environmental Microbiology, 82(8), 2506–2515.

6. Yan, S., Wang, S., Wei, G. & Zhang, K. (2015). Investigation of the main parameters during the fermentation of Chinese Luzhou – flavour liquor. Journal of the Institute of Brewing, 121(1), 145–154.

7. Tao, Y., Wang, X., Li, X., Wei, N., Jin, H., Xu, Z., Tang, Q. & Zhu, X. (2017). The functional potential and active populations of the pit mud microbiome for the production of Chinese strong – flavour liquor. Microbial Biotechnology, 10(6), 1603–1615.

8. Fan, W. & Qian, M. C. (2010). Identification of aroma compounds in Chinese ‘Yanghe Daqu’ liquor by normal phase chromatography fractionation followed by gas chromatography[sol] olfactometry. Flavour & Fragrance Journal, 21(2), 333–342.

9. Zheng, X. W. & Han, B. Z. (2016). Baijiu (白酒), Chinese liquor: History, classification and manufacture. Journal of Ethnic Foods, 3(1), 19–25.

10. Xu, Y., Wang, D., Fan, W. L., Mu, X. Q. & Chen, J. (2010). Traditional Chinese biotechnology. Advances in Biochemical Engineering/biotechnology, 122(189–233.

11. Zheng, J., Liang, R., Zhang, L., Wu, C., Zhou, R. & Liao, X. (2013). Characterization of microbial communities in strong aromatic liquor fermentation pit muds of different ages assessed by combined DGGE and PLFA analyses. Food Research International, 54(1), 660–666.

12. Wu, Z. Y., Zhang, W. X., Zhang, Q. S., Cheng, H., Rong, W. & Liu, Z. H. (2012). Developing new sacchariferous starters for liquor production based on functional strains isolated from the pits of several famous Luzhou-flavor liquor brewers. Journal of the Institute of Brewing, 115(2), 111–115.

13. Yue, Y., Zhang, W., Liu, X., Hu, C. & Zhang, S. (2007). Isolation and identification of facultative anaerobes in the pit mud of Chinese Luzhou-flavor liquor. Microbiology, 34(2), 251–255.

14. Amann, R. I., Ludwig, W. & Schleifer, K. H. (1995). Phylogenetic identification and in situ detection of individual microbial cells without cultivation. Microbiol Rev, 59(1), 143–169.

15. Zhao, J. S., Zheng, J., Zhou, R. Q. & Shi, B. (2012). Microbial community structure of pit mud in a Chinese strong aromatic liquor fermentation pit. Journal of the Institute of Brewing, 118(4), 356–360.

16. Hong, X., Jing, C., Lin, L., Wu, H., Tan, H., Xie, G., Qian, X., Zou, H., Yu, W. & Lan, W. (2016). Metagenomic sequencing reveals the relationship between microbiota composition and quality of Chinese Rice Wine. Scientific Reports, 6, 26621.

17. Glenn, T. C. (2011). Field guide to next – generation DNA sequencers. Molecular Ecology Resources, 11 (5), 759–769.

18. Yarza, P., Yilmaz, P., Pruesse, E., Glöckner, F. O., Ludwig, W., Schleifer, K. H., Whitman, W. B., Euzéby, J., Amann, R. & Rosselló-Móra, R. (2014). Uniting the classification of cultured and uncultured bacteria and archaea using 16S rRNA gene sequences. Nature Reviews Microbiology, 12(9), 635–645.

19. Liang, H., Li, W., Luo, Q., Liu, C., Wu, Z. & Zhang, W. (2015). Analysis of the bacterial community in aged and aging pit mud of Chinese Luzhou-flavour liquor by combined PCR-DGGE and quantitative PCR assay. Journal of the Science of Food & Agriculture, 95(13), 2729–2735.

20. Zheng, Q., Lin, B., Wang, Y., Zhang, Q., He, X., Yang, P., Zhou, J., Guan, X. & Huang, X. (2015). Proteomic and high-throughput analysis of protein expression and microbial diversity of microbes from 30- and 300-year pit muds of Chinese Luzhou-flavor liquor. Food Research International, 75, 305–314.

21. Shin, J., Lee, S., Go, M. J., Sang, Y. L., Sun, C. K., Lee, C. H. & Cho, B. K. (2016). Analysis of the mouse gut microbiome using full-length 16S rRNA amplicon sequencing. Scientific Reports, 6, 29681.

22. Caporaso, J. G., Kuczynski, J., Stombaugh, J., Bittinger, K., Bushman, F. D., Costello, E. K., Fierer, N., Peña, A. G., Goodrich, J. K. & Gordon, J. I. (2010). QIIME allows analysis of high-throughput community sequencing data. Nat Methods., 335–336.

23. Edgar, R. C. (2010). Search and clustering orders of magnitude faster than BLAST. Bioinformatics, 26(19), 2460–2461.

24. Sradnick, A., Murugan, R., Oltmanns, M., Raupp, J. & Joergensen, R. G. (2013). Changes in functional diversity of the soil microbial community in a heterogeneous sandy soil after long-term fertilization with cattle manure and mineral fertilizer. Applied Soil Ecology, 63(63), 23–28.

25. Lennon, J. T., Aanderud, Z. T., Lehmkuhl, B. K. & Jr, S. D. (2012). Mapping the niche space of soil microorganisms using taxonomy and traits. Ecology, 93(8), 1867–1879.

26. Roelke, D. L., Kokkoris, G. D., Spatharis, S. & Dimitrakopoulos, P. G. (2011). Analyzing the (mis)behavior of Shannon index in eutrophication studies using field and simulated phytoplankton assemblages. Ecological Indicators, 11 (2), 697–703.

27. Wang, Y., Sheng, H. F., He, Y., Wu, J. Y., Jiang, Y. X., Tam, N. F. & Zhou, H. W. (2012). Comparison of the levels of bacterial diversity in freshwater, intertidal wetland, and marine sediments by using millions of illumina tags. Applied & Environmental Microbiology, 78(23), 8264–8271.

28. Ding, X., Wu, C., Huang, J. & Zhou, R. (2015). Interphase microbial community characteristics in the fermentation cellar of Chinese Luzhou -flavor liquor determined by PLFA and DGGE profiles. Food Research International, 72, 16–24.

29. Guo, M. Y., Huo, D. Q., Ghai, R., Rodriguezvalera, F., Shen, C. H., Zhang, N., Zhang, S. Y. & Hou, C. J. (2014). Metagenomics of ancient fermentation pits used for the production of chinese strong-aroma liquor. Genome Announcements, 2(5), 32–45.

30. Barberan, A., Bates, S. T., Casamayor, E. O. & Fierer, N. (2012). Using network analysis to explore co-occurrence patterns in soil microbial communities. ISME Journal, 6(2), 343–351.

31. Lupatini, M.; Suleiman, A. K. A.; Jacques, R. J. S.; Antoniolli, Z. I.; De, A.; Ferreira, S.; Kuramae, E. E.; & Roesch, L. F. W. (2014). Network topology reveals high connectance levels and few key microbial genera within soils. Frontiers in Environmental Science, 2(10), 1–11.

32. Peura, S., Bertilsson, S., Jones, R. I. & Eiler, A. (2015). Resistant microbial cooccurrence patterns inferred by network topology. Applied & Environmental Microbiology, 81(6), 2090–2097.

33. Vanwonterghem, I., Jensen, P. D., Dennis, P. G., Hugenholtz, P., Rabaey, K. & Tyson, G. W. (2014). Deterministic processes guide long-term synchronised population dynamics in replicate anaerobic digesters. Isme Journal, 8(10), 2015–2028.

34. Ding, X. F., Wu, C. D., Zhang, L. Q., Zheng, J. & Zhou, R. Q. (2014). Characterization of eubacterial and archaeal community diversity in the pit mud of Chinese Luzhou-flavor liquor by nested PCR-DGGE. World Journal of Microbiology & Biotechnology, 30(2), 605–612.

35. Breitenstein, A., Wiegel, J., Haertig, C., Weiss, N., Andreesen, J. R. & Lechner, U. (2002). Reclassification of *Clostridium hydroxybenzoicum* as *Sedimentibacter hydroxybenzoicus* gen. nov., comb. nov., and description of *Sedimentibacter saalensis* sp. nov. International Journal of Systematic & Evolutionary Microbiology, 52, 801–807.

36. Gies, E. A., Konwar, K. M., Beatty, J. T. & Hallam, S. J. (2014). Illuminating microbial dark matter in meromictic Sakinaw Lake. Applied & Environmental Microbiology, 80(21), 6807–6818.

37. Zhang, C. & Liu, X. X. (2004). *Syntrophomonas curvata* sp. nov., an anaerobe that degrades fatty acids in co-culture with methanogens. International Journal of Systematic & Evolutionary Microbiology, 54(3), 969–973.

38. López, N. I., Pettinari, M. J., Stackebrandt, E., Tribelli, P. M., Põtter, M., Steinbüchel, A. & Méndez, B. S. (2009). *Pseudomonas extremaustralis* sp. nov., a poly(3-hydroxybutyrate) producer isolated from an Antarctic environment. Current Microbiology, 59(5), 514–519.

39. Catone, M. V., Ruiz, J. A., Castellanos, M., Segura, D., Espin, G. & López, N. I. (2014). High polyhydroxy butyrate production in *Pseudomonas extremaustralis* is associated with differential expression of horizontally acquired and core genome polyhydroxyalkanoate synthase genes. Plos One, 9(6), e98873.

40. Collins, M. D., Lawson, P. A., Willems, A., Cordoba, J. J., Fernandez-Garayzabal, J., Garcia, P., Cai, J., Hippe, H. & Farrow, J. A. (1994). The phylogeny of the genus Clostridium: proposal of five new genera and eleven new species combinations. Int J Syst Bacteriol, 44(4), 812–826.

41. Louis, P., & Flint, H. J. (2009). Diversity, metabolism and microbial ecology of butyrate-producing bacteria from the human large intestine. FEMS Microbiology Letters, 294, 1–8.

42. Wu, C., Liu, X. & Dong, X. (2006). *Syntrophomonas erecta* subsp. *sporosyntropha* subsp. nov., a spore-forming bacterium that degrades short chain fatty acids in co-culture with methanogens. Systematic & Applied Microbiology, 29(6), 457–462.

43. Alsaker, K. V.; Paredes, C.; Papoutsakis, E. T. (2010) Metabolite stress and tolerance in the production of biofuels and chemicals: Gene-expression-based systems analysis of butanol, butyrate, and acetate stresses in the anaerobe Clostridium acetobutylicum. Biotechnology and Bioengineering, 105, 1131–1147.

